# Asi1 regulates the distribution of proteins at the inner nuclear membrane in *Saccharomyces cerevisiae*

**DOI:** 10.1101/442970

**Authors:** Christine J. Smoyer, Sarah E. Smith, Scott McCroskey, Jay R. Unruh, Sue L. Jaspersen

## Abstract

Inner nuclear membrane (INM) protein composition regulates nuclear function, affecting processes such as gene expression, chromosome organization, nuclear shape and stability. Mechanisms that drive changes in the INM proteome are poorly understood in part because it is difficult to definitively assay INM composition rigorously and systematically. Using a split-GFP complementation system to detect INM access, we examined the distribution of all C-terminally tagged *Saccharomyces cerevisiae* membrane proteins in wild-type cells and in mutants affecting protein quality control pathways, such as INM-associated degradation (INMAD), ER-associated degradation (ERAD) and vacuolar proteolysis. Deletion of the E3 ligase Asi1 had the most pronounced effect on the INM compared to mutants in vacuolar or ER-associated degradation pathways, consistent with a role for Asi1 in the INMAD pathway. Our data suggests that Asi1 not only removes mis-targeted proteins at the INM, but it also controls the levels and distribution of native INM components, such as the membrane nucleoporin Pom33. Interestingly, loss of Asi1 does not affect Pom33 protein levels but instead alters Pom33 distribution in the NE through Pom33 ubiquitination, which drives INM redistribution. Taken together, our data demonstrate that the Asi1 E3 ligase has a novel function in INM protein regulation in addition to protein turnover.

## Introduction

The nucleus is the defining feature of eukaryotic cells. Successful propagation of nuclei, and the genome contained within their walls, is vital for an organism’s survival. Although the double lipid bilayer that forms the nuclear envelope (NE) is often viewed as a fortress, the inner and outer nuclear membranes (INM and ONM) are highly dynamic structures that undergo changes in structure and composition throughout development and differentiation, in mitosis and meiosis, and in diseased and dying cells (Chen *et al.* 2014; Dauer and Worman 2009; Gordon *et al.* 2014). Lobulated nuclei with aberrant membranes and abnormal chromosome configurations are used to grade many tumors; however, similar changes in nuclear shape and genome organization also occur during the maturation of normal cell types, most notably during hematopoiesis, suggesting a complexity at the NE that we are just beginning to understand (Skinner and Johnson 2017). As many of the unique properties of the NE, such as its mechanical stiffness, distinctive lipid composition and chromosome organization, are attributed to the INM, understanding the composition, function and regulation of the INM is a key problem in cell biology.

In all eukaryotes, the INM and ONM are joined together at many spots where nuclear pore complexes (NPCs) reside. NPCs form the first regulator of INM composition by controlling the passage of proteins and other macromolecules into and out of the nucleus. INM proteins travel through central or lateral channels of the NPC to reach the INM (reviewed in (Katta *et al.* 2014; Ungricht and Kutay 2017)). Most do not have any targeting sequence and reach the INM by diffusion; their retention at the INM occurs through binding to nuclear or NE-associated proteins such as lamins, NPCs or chromatin (Antonin *et al.* 2011; Furukawa *et al.* 1998; Ungricht *et al.* 2015; Ungricht and Kutay 2015; Wu *et al.* 2002). A small subset of proteins is targeted to the INM by a specific sequence motif, which is recognized by the nuclear translocation machinery (Gardner *et al.* 2011; King *et al.* 2006; Lusk *et al.* 2007; Tapley *et al.* 2011; Turgay *et al.* 2010). Additional mechanisms of INM transport have also been proposed that bypass the NPC and rely on the budding and fusion of vesicles from ONM to INM (Mettenleiter 2016; Speese *et al.* 2012). While many mechanistic details regarding INM transport are still poorly understood, it seems clear that the INM has a distinct composition from the ONM, which is contiguous with the endoplasmic reticulum (ER).

In the ER, misfolded or damaged proteins are targeted for degradation by the ER-associated degradation pathway (ERAD) as part of the cell’s surveillance system to prevent the formation of non-functional complexes or aggregates of defective protein (Zattas and Hochstrasser 2015). The conserved E3 ligases Doa10/MARCH6/TEB4 and Hrd1/SYVN1 recognize lesions in the cytosolic or luminal/membrane regions of ER proteins, respectively, resulting in ubiquitination and retro-translocation of defective proteins back into the cytoplasm for destruction by the 26S proteasome (Zattas and Hochstrasser 2015). Although ERAD likely ensures that ONM proteins are functional and present in the correct stoichiometry, it is unclear if this pathway operates at the INM due to its separation from the ONM/ER by NPCs (Boban and Foisner 2016). Instead, an INM-associated degradation (INMAD) has been proposed to remove mistargeted proteins from the INM through ubiquitin-mediated proteolysis (Foresti *et al.* 2014; Khmelinskii *et al.* 2014). Three putative E3 ligases have been implicated in INMAD in yeast (Doa10, Asi1 and Asi3) and a handful of substrates have been identified based on increased whole cell protein levels in cells lacking the ligases. Interestingly, these INMAD substrates are not INM components but are proteins mistargeted to the INM, such as a mutant version of the Sec61 translocon, vacuolar transport complex subunits and multiple plasma membrane transporters (Foresti *et al.* 2014; Khmelinskii *et al.* 2014). The mechanisms by which INMAD distinguishes between ‘foreign’ and ‘resident’ membrane proteins, and whether INMAD targets damaged or misfolded INM components through a pathway similar to ERAD remains unknown. Examination of protein stability in rats using isotope labeling suggested that NPCs and INM proteins such as lamins are extremely long-lived, leading to the general idea that the INM is stable, with little protein turnover (Savas *et al.* 2012). One important exception occurs under nutrient deprivation when non-essential sections of the entire nucleus, including the INM, are pinched off into the vacuole (the yeast lysosome) and degraded, a process known as piecemeal nuclear autophagy (PMNA) (Adnyana *et al.* 2000; Do *et al.* 2003; Millen *et al.* 2009; Roberts *et al.* 2003). PMNA has not yet been identified in higher eukaryotes, but autophagy of NE proteins in mammalian cells occurs (Dou *et al.* 2015; Park *et al.* 2009) and is linked to both cancer and aging (Martinez-Lopez *et al.* 2015; White *et al.* 2015). However, a role for INMAD outside of yeast has not been reported.

We previously described a method using split-GFP that allowed us to systematically and unequivocally assay the ability of budding yeast membrane proteins to access the INM (Smoyer *et al.* 2016). Unlike biochemical methods for studying INM composition that depend on in silico subtraction or comparative analysis of nuclear and microsomal membrane samples (an ER derived fraction formed in vitro), our assay is specific for the INM pool of protein, allowing for analysis of proteins that have dual functions. The assay also discriminates the INM from ONM using endogenously expressed proteins that serve as the sole copy in the cell and can be used in live cells. To test the contribution of INMAD on INM composition, we compared the distribution of proteins using split-GFP in wild-type cells to that of cells lacking *ASI1*. Comparison of INM composition in cells lacking other protein quality control components such as the INMAD/ERAD E3 ligase Doa10, the ERAD E3 ligase Hrd1, the E2 ubiquitin conjugating enzyme Ubc7 and the vacuolar peptidase Pep4, revealed 74 proteins whose INM levels increased specifically in cells lacking *ASI1*, suggesting direct or indirect regulation by Asi1. In *asi1∆*, we also observed an increase in the size and frequency of INM puncta containing Pom33/Tts1/TMEM33, a conserved NPC-localized protein that plays a role in NPC distribution, biogenesis and/or stability (Chadrin *et al.* 2010; Urade *et al.* 2014; Zhang and Oliferenko 2014). We provide evidence that Pom33 is directly ubiquitinated by Asi1; however, ubiquitination of Pom33 does not lead to turnover, but instead contributes to proper Pom33 INM distribution.

## Results

### Screen for INM changes in protein quality control mutants

To detect if proteins are able to access the INM, we express a soluble nuclear protein at high levels fused to half of split-GFP (GFP_11_-mCherry-Pus1); this can reconstitute a working GFP when a protein fused to GFP_1-10_ localizes to the same compartment (Figure 1A). Previously, we showed that 312 of 1010 C-terminally tagged membrane or predicted membrane proteins in yeast have access to the INM using the split-GFP assay (Smoyer *et al.* 2016). To determine how the INM proteome is altered by removal of quality control systems, we examined INM localization of the same library in cells lacking *ASI1* (INMAD), *DOA10* (INMAD/ERAD), *HRD1* (ERAD), *UBC7* (INMAD/ERAD and other pathways) and *PEP4* (vacuole). Wild-type and mutant strains were screened using a high-throughput 96-well plate imaging format, and images were quantitatively assessed for INM signal using an automated image analysis pipeline. Although it is unlikely that the amount of nuclear reporter is limiting as it is present in high copy, levels of GFP_11_-mCherry-Pus1 could affect the amount of 488 fluorescence visualized, so we first normalized to mCherry levels on a cell by cell basis. Next, we averaged split-GFP signal for each protein/mutant combination to eliminate possible cell cycle/cell growth artifacts and then used this value to determine the final intensity ratio of wild-type and each mutant (Table S1).

**Figure 1.**
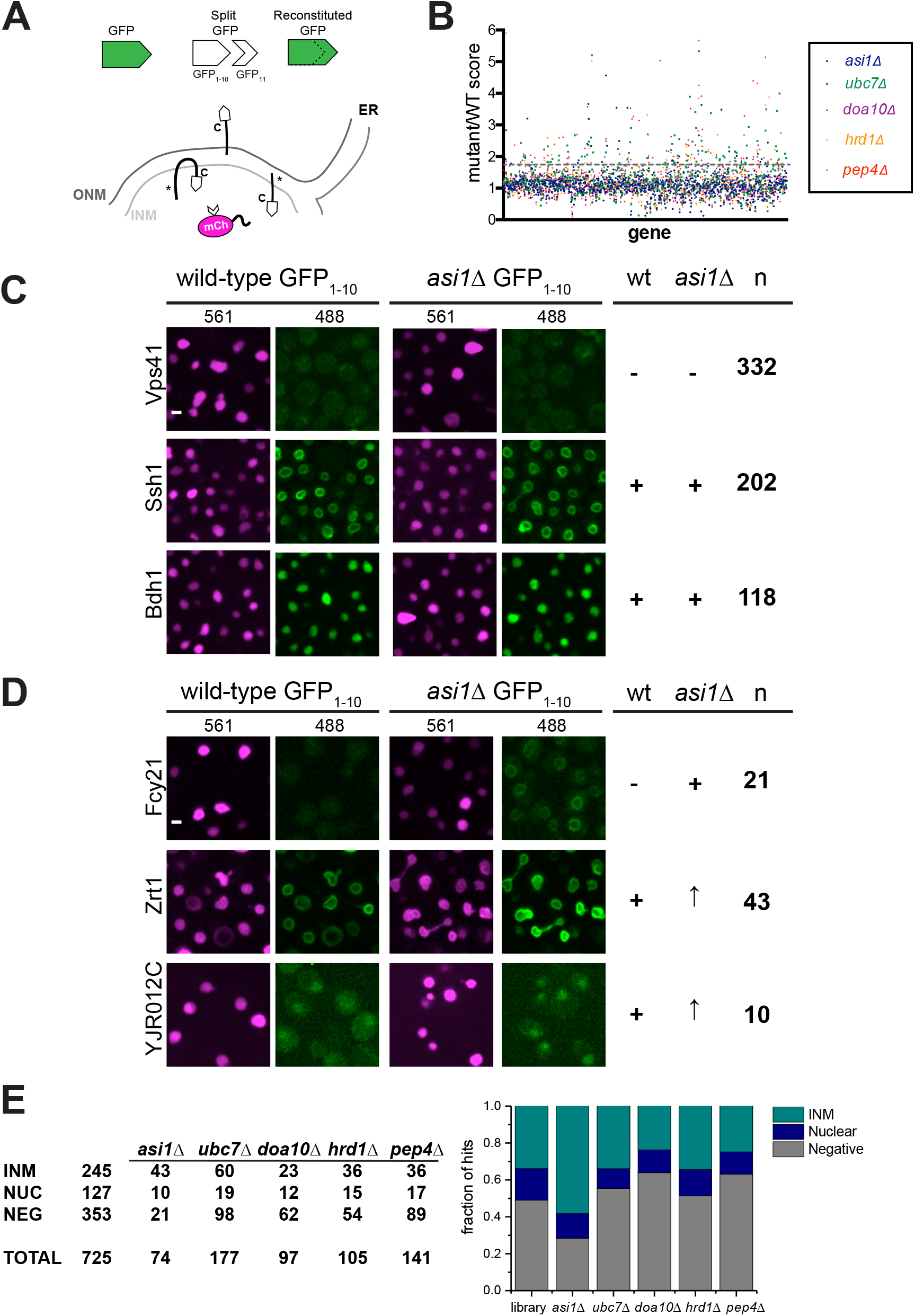
Mutation of *ASI1* alters split-GFP signal of a subset of INM proteins. A. Schematic showing split-GFP and its use to detect proteins that can access the INM (denoted with asterisk). B. Plot of the split-GFP mutant/wild-type ratios for each gene reported from 96-well plate imaging analysis of membrane proteins in *asi1∆*, *ubc7∆*, *doa10∆*, *hrd1∆* and *pep4∆*. Dashed line represents the 1.4 cut-off. C. Representative images of proteins that were categorized previously as negative or positive (INM or soluble) that remained unchanged in *asi1∆*, based on *asi1∆*/wild-type ratio of 1.39 or lower. D. Signal of proteins that changed in *asi1∆* largely fell into 3 categories: proteins that were previously negative, and proteins that were positive (INM or soluble) that increased in *asi1∆*. The number of proteins assigned to each category for B and C is on the right, based on *asi1∆*/wild-type ratio of 1.4 or higher. E. Left, table summary of starting library localization based on wild-type results in the first column, and the number represented in each list of mutant hits based on a ratio of 1.4 or higher, mutant/wild-type. Right, graphical representation of corresponding table. Bars, 2 μm.

Manual inspection of images in cells lacking *ASI1* compared to wild-type showed that only 725 of the 833 genes tested in the mutant could be reliably scored. Most (84/105) of the 105 false positives (Table S2) correspond to proteins absent from the INM of both wild-type and *asi1∆*; for the remainder (21/105), we suspect that low expression levels and higher levels of autofluorescence resulted in ambiguous scoring, so these were eliminated from our library and were not included in analysis of any mutants. The mating pheromone receptor Ste2 was also a false positive due to the MATa mating type of strains used for mutant analysis versus the MATα strain we used for wild-type (Figure S1). If we examined Ste2-GFP_1-10_ in a wild-type MATa strain, it localizes to the INM like the mutants (Figure S1). No other INM components were obviously affected by mating type.

Our analysis showed that the distribution of most proteins was unaffected in the deletions, resulting in ratio values around 1 (Figure 1B-C). Using a cut-off of 1.4, which is slightly less than one standard deviation above the mean for most samples, we found that 10-26% of proteins showed an increase above wild-type. This included increased levels for 74/725 proteins in *asi1∆*, 177/672 in *ubc7∆*, 97/705 in *doa10∆*, 105/707 in *hrd1∆* and 141/686 in *pep4∆* (Figure 1E). These hits could be further subdivided into three categories based on their localization in wild-type. For example, 21 proteins present at the INM in *asi1∆* did not localize to the INM in wild-type, 10 were soluble and nucleoplasmic and 43 were at the INM but increased in cells lacking *ASI1* (Figure 1D-E). This over-representation of INM components in *asi1∆* (58%) compared to other mutants (34%) strongly suggests that Asi1 affects native INM components. Consistent with this idea, the well-characterized INM components Pom33, Per33, Asi2 and Heh2 were identified as hits in our *asi1∆* screen but not in other mutants (Table S1 and S3), as discussed below.

In each mutant analyzed, we identified increased INM access of a unique combination of split-GFPs (Table S3). Cells lacking the E2 ubiquitin-conjugating enzyme Ubc7 partially overlapped with the E3 ligases, consistent with the idea that Ubc7 plays a role in both ERAD and INMAD. The partial overlap between *doa10∆* and *hrd1∆* or *asi1∆* mutants (Figure S2A) is similar to previous reports suggesting that Doa10 also plays a role in both pathways (Foresti *et al.* 2014; Khmelinskii *et al.* 2014). The limited overlap between our split-GFP library and the tandem timer library used by Khmelinskii et al only allowed us to test 12 of 20 previously reported Asi1 substrates (Khmelinskii *et al.* 2014). From this, we confirmed increased INM localization in eight of the twelve, including the vacuolar transferase complex subunit Vtc1 that is thought to be mistargeted to the INM due to protein tagging (Khmelinskii *et al.* 2014), the Rab GTPase interacting protein Yip4, the plasma membrane transporter Zrt2, inositolphosphotransferase Ipt1, and Irc23, a protein of unknown function that is linked to DNA damage (Figure S2B). We also saw similar effects for *ubc7∆*, but interestingly, our screen showed little or no overlap in *doa10∆* and *hrd1∆* substrates (Figure S2B) (Foresti *et al.* 2014; Khmelinskii *et al.* 2014). It is unknown if protein levels and/or INM localization is a direct or indirect consequence of the Asi1 deletion; therefore, we performed more detailed analysis on a subset of mutants and proteins.

### Pom33 distribution is altered in cells lacking ASI1

In addition to a change in protein levels, one of the most interesting phenotypes that we observed in INM quality control mutants was the appearance of puncta—increased levels of INM protein at one or more NE locations in the presence of background fluorescence throughout the membrane. A striking example is shown in Figure 2A for the paralogs, Pom33 and Per33, in cells lacking *ASI1*. The INM component Heh2 and the NPC pore membrane protein Pom34 did not form puncta in any mutant analyzed (Figure 2A & S3), suggesting that puncta formation is specific to Pom33 and Per33 and not a general feature of INM or NPC pore components in *asi1∆*. Pom34 was identified as a potential Ubc7-dependent, Hrd1- and Doa10-independent target in a previous proteomic screen for Asi1 targets (Foresti *et al.* 2014) and in our analysis, it did exhibit a slight ratio increase for *asi1∆* (1.27) and *ubc7∆* (1.35). Unlike the puncta formation we previously observed for a number of proteins using split-GFP (Smoyer *et al.* 2016), the puncta formed by Pom33 were specifically linked to INM quality control pathways, forming in mutant cells lacking *ASI1*, *ASI2*, *UBC7* and to a lesser extent *ASI3* (Figure 2A, 2C & S3). Per33 puncta also formed in ERAD mutants (Figure S3) and were not studied further. Quantitation of Pom33 at the INM showed that there was little change in intensity or total Pom33 levels in wild-type and mutants (Figure 2A & B), but the frequency of puncta more than doubled in *asi1∆* and *asi2∆* cells (Figure 2C). These structures were also considerably larger than in wild-type cells (Figure 2D), suggesting that Asi1 and Asi2 affect Pom33 distribution at the INM.

**Figure 2.**
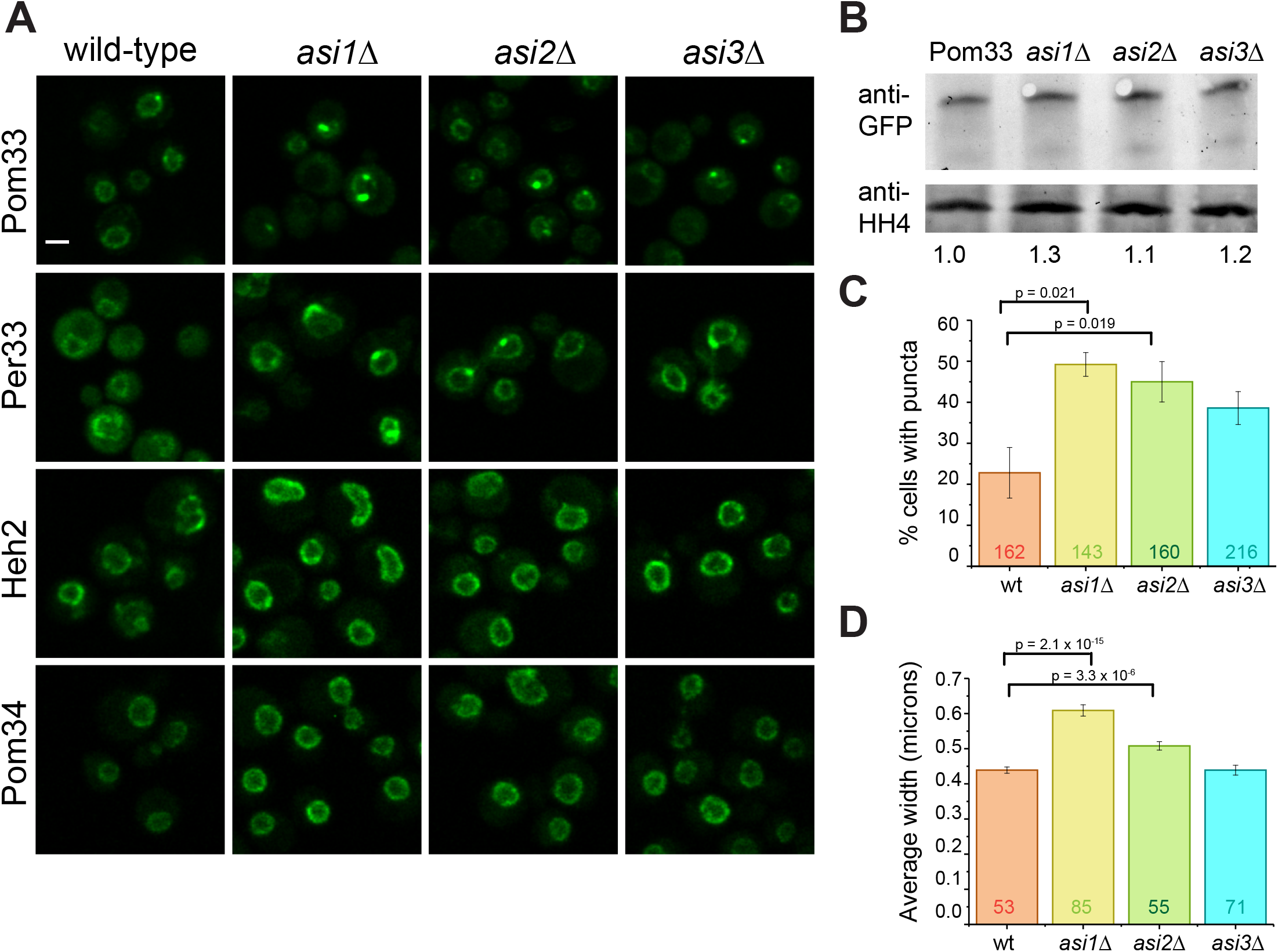
Pom33 distribution is altered in cells lacking *ASI1*, *ASI2* or *ASI3*. A. Comparison of GFP_1-10_ signal for Pom33, Per33, Heh2 and Pom34 in wild-type and cells lacking *ASI1*, *ASI2* or *ASI3*. B. Quantitation of total protein levels for Pom33 in wild-type, *asi1∆*, *asi2∆* and *asi3∆*. Pom33-GFP_1-10_ levels were first normalized to histone H4 signal and the mutant levels each compared to wild-type; ratios set at bottom. C. The frequency of Pom33-GFP1-10 puncta increases in mutants of *ASI1* and *ASI2*. Total cells counted depicted on graph. D. The average puncta width measured by full width at half maximum (FWHM) of fluorescence intensity at the puncta. Total puncta measured depicted on graph. Bar, 2 μm. p values were determined by Students t-test. Error bars equal SEM.

### Ubiquitination of Pom33 regulates INM distribution but not degradation

To understand how Asi1 and Asi2 affect Pom33 distribution at the INM, we considered the possibility that Pom33 is a target of the ubiquitin ligase activity of Asi1 and is ubiquitinated in vivo. Ubiquitinated proteins were immunoprecipitated from lysates containing Pom33 and Pom33-GFP_1-10_, with and without *ASI1* with an anti-ubiquitin antibody. As a positive control, we immunoprecipitated an N-terminal fusion of ubiquitin to Pom33-GFP_1-10_ (Ub-Pom33). We analyzed these immunoprecipitates by western blotting with an anti-GFP antibody. Although a ladder of bands was not present to indicate poly-ubiquitination, we enriched for Pom33-GFP_1-10_ from wild-type but not *asi1∆* cells in these experiments (Figure 3A-B), suggesting that Pom33 is ubiquitinated in an Asi1-dependent manner.

**Figure 3.**
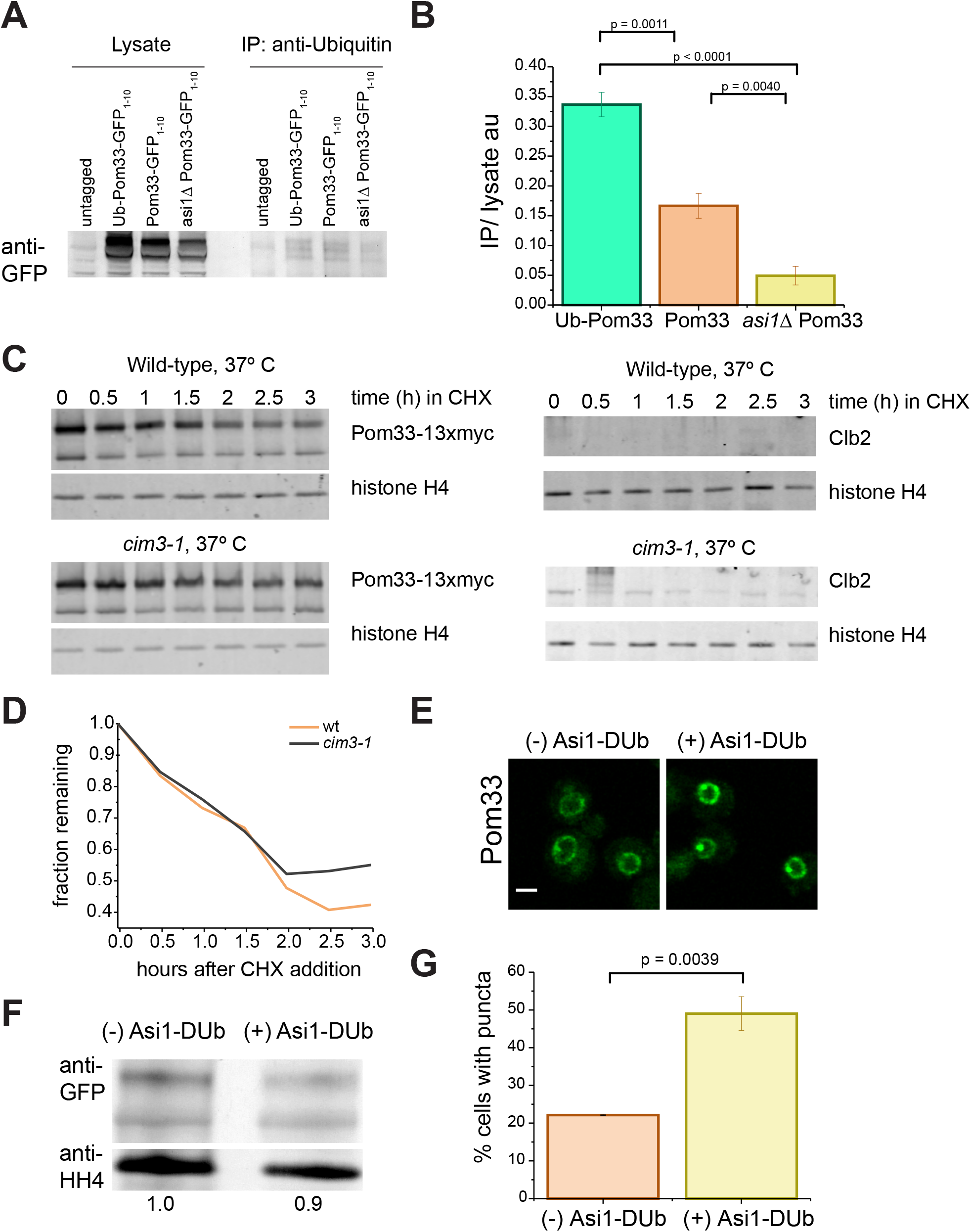
Asi1 dependent ubiquitination of Pom33 does not lead to its degradation by the proteasome. A. Immunoprecipitation (IP) of an untagged control strain, Ubiquitin-Pom33-GFP_1-10_, Pom33-GFP_1-10_ and *asi1∆* Pom33-GFP_1-10_ using an anti-Ubiquitin antibody, followed by immunoblot analysis using an anti-GFP antibody. B. Quantitation of signal of Pom33-GFP_1-10_ in IP over signal in lysate, as determined by four separate IP experiments. Error bars equal SEM. C. Time course of total levels of Pom33-13xmyc in wild-type and *cim3-1* at 37°. Cells were shifted to the nonpermissive temperature for 45 min before addition of cycloheximide. Results of western blot analysis at the given time points for Pom33-13xmyc on the left, and Clb2 on the right. D. Ratio of Pom33-GFP_1-10_ protein remaining as determined by western blot. E. Pom33–GFP_1-10_ with and without Asi1-DUb expressed. F. Total levels of Pom33-GFP_1-10_ with and without Asi1-DUb expressed. G. Frequency of Pom33-GFP_1-10_ puncta, DUb off: n = 95 cells, DUb on: n = 96 cells. Bar, 2 μm. P values were determined by Students t-test.

Total Pom33 proteins levels do not change in *asi1∆* (Figure 2B) and it does not appear to be poly-ubiquitinated (Figure 3A), suggesting Pom33 is not actively degraded. To formally test whether Pom33 is degraded by the 26S proteasome, we examined the stability of Pom33-13xmyc in wild-type cells compared to *cim3-1,* a mutant that disrupts the proteasome lid component Rpt6 (Ghislain *et al.* 1993; Schork *et al.* 1995), in a cycloheximide chase experiment at the nonpermissive temperature of 37°C. After wild-type and *cim3-1* cells were shifted to 37°C for 45 min, cycloheximide was added to inhibit protein synthesis, and Pom33-13xmyc protein levels were monitored by western blot analysis at the indicated times (Figure 3C-D). Clb2, a cell cycle protein known to be degraded by the proteasome (Deshaies 1997; Seufert *et al.* 1995), served as a control that showed stabilization in *cim3-1* (Figure 3C, right), while histone H4 was a loading control since its levels and stability are unaffected by temperature, cycloheximide or proteasome inhibition. Pom33-13xmyc stability was unaffected by *cim3-1* (Figure 3D), suggesting that Pom33 is not a proteasome target.

If Pom33 is ubiquitinated, but not degraded, what role does Asi1-dependent ubiquitination play in regulation and function of Pom33? One attractive idea is that Asi1-dependent ubiquitination may modulate Pom33 INM distribution, similar to the way that ubiquitination of proteins is involved in different processes such as nucleocytoplasmic trafficking. Examples of this include p53, in which mono-ubiquitination is thought to expose a nuclear export signal (Li *et al.* 2003; LOHRUM *et al.* 2001; Nie *et al.* 2007) and cytidydyltransferase (CCTalpha) where mono-ubiquitination blocks its nuclear localization signal (Chen and Mallampalli 2009). To test this idea, we employed an inducible version of Asi1 fused to the deubiquitinating domain of Herpes Virus UL36 (Asi1-DUb) (Macdonald *et al.* 2012; Macdonald *et al.* 2017). This Asi1-DUb is designed to deubiquitinate Asi1 targets as Asi1 acts on them. Although Asi1-DUb had little effect on total Pom33 protein levels, its induction resulted in increased frequency and extent of Pom33-GFP_1-10_ puncta formation at the INM (Figure 3E-G), a phenotype very similar to the *asi1∆*. The appearance of the Pom33 puncta upon induction of the Asi1-DUb suggest that their formation is a direct consequence of deubiquitination and supports the idea that Asi1-dependent ubiquitination of Pom33 regulates its INM distribution.

If ubiquitination of Pom33 contributes to its normal INM distribution, then constitutive ubiquitination by fusion of ubiquitin to Pom33 might rescue the puncta phenotype seen in *asi1∆* mutants. To test this, we first measured the INM intensity, using split-GFP signal, of Pom33-GFP_1-10_ and Ub-Pom33-GFP_1-10_ in wild-type and *asi1∆* cells by split-GFP levels, finding that Ub-Pom33-GFP_1-10_ partially recused overall INM intensity in *asi1∆* (Figure 4A-B). Next, we compared the frequency and size of puncta in wild-type and *asi1∆* containing Pom33-GFP_1-10_ or Ub-Pom33-GFP_1-10_ expressed in single copy at the *URA3* locus as the sole copy of *POM33* in the cell. As expected, the frequency and size of Pom33-GFP_1-10_ puncta increased in *asi1∆* compared to wild-type (Figure 4A-C). However, expression of Ub-Pom33-GFP_1-10_ rescued the frequency and size phenotypes in *asi1∆* cells, with fewer puncta (Figure 4A-C) as well as smaller puncta size (Figure 4C, bottom). Therefore, attaching ubiquitin to Pom33 bypasses the requirement for Asi1 for INM distribution, strongly suggesting Asi1 directly modifies Pom33. Because modification of N-terminal residues can affect protein stability through the N-end rule pathway (Tasaki *et al.* 2012; Varshavsky 1992), we examined expression levels by western blotting. This analysis showed that Pom33-GFP_1-10_ and Ub-Pom33-GFP_1-10_ were present at similar levels in wild-type cells (Figure 4D).

**Figure 4.**
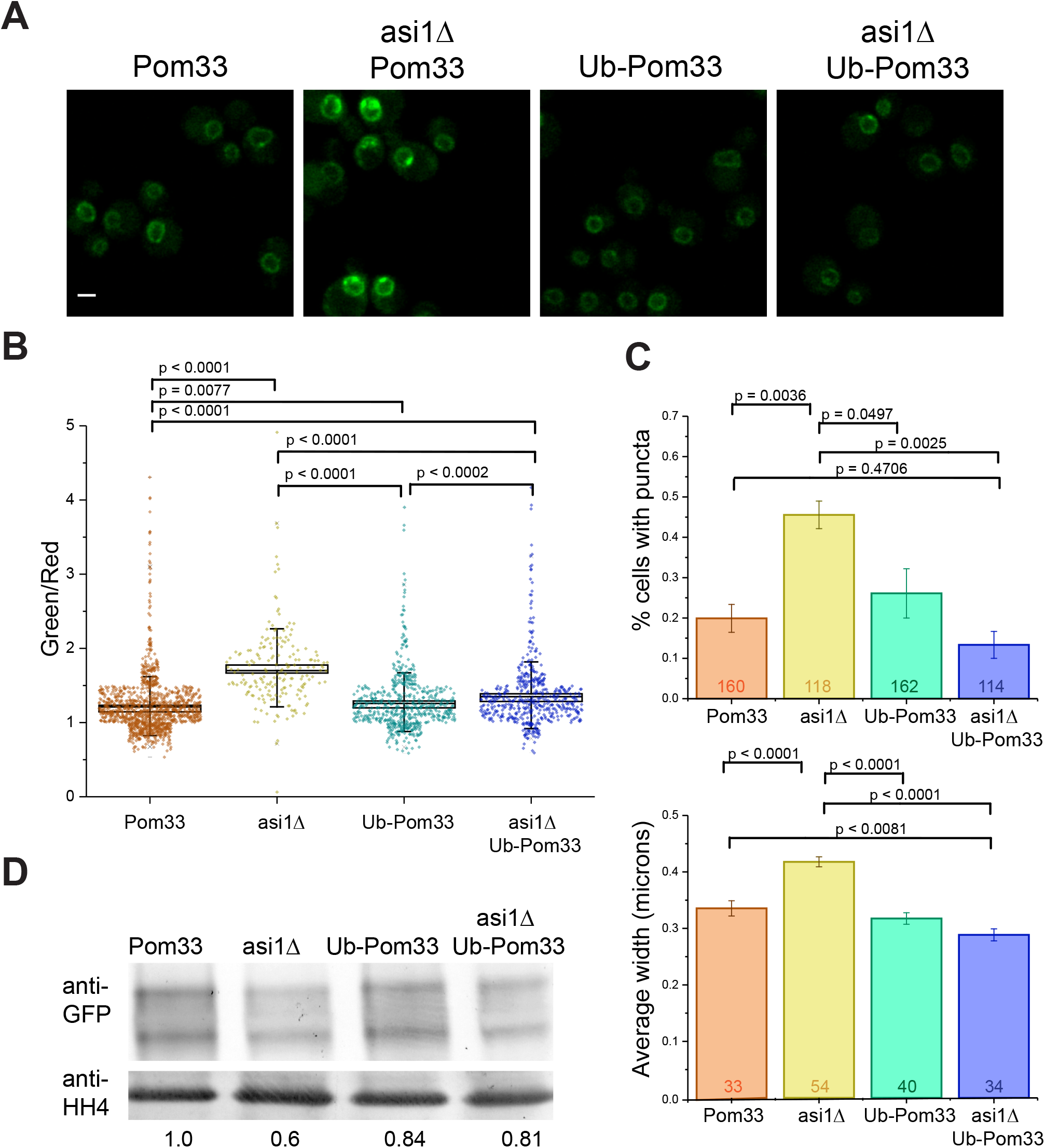
Expression of Ub-Pom33-GFP_1-10_ reduces puncta frequency and size in *asi1∆*. A. Example images of Pom33-GFP_1-10_, *asi1∆* Pom33-GFP_1-10_, Ub-Pom33-GFP_1-10_ and *asi1∆* Ub-Pom33-GFP_1-10_. B. Quantification of strains in A; intensities measured as a ratio of GFP^1-10^ (488) signal over Pus1 (561) signal. C. For strains depicted in A, the frequency of Pom33-GFP_1-10_ puncta was counted (top) and the average puncta width measured by (FWHM) of fluorescence intensity at the puncta (bottom). Total n depicted on graphs. Error bars equal SEM. D. Quantitation of total protein levels for Pom33-GFP_1-10_, *asi1∆* Pom33-GFP_1-10_, Ub-Pom33-GFP_1-10_ and *asi1∆* Ub-Pom33-GFP_1-10_ by western blot. Levels were first normalized to histone H4 signal and the levels of each compared to wild-type on left; ratios set at bottom. Bar, 2 μm.

### Pom33 INM puncta contain NPC components

Punctate distribution at the NE has been previously reported for spindle pole body (SPB), nuclear-vacuolar junction (NVJ), intranuclear quality control (INQ) and storage for incomplete NPC (SINC) components: Spc42, Nvj1, Cmr1 and Chm7, respectively (Gallina *et al.* 2015; Huh *et al.* 2003; Webster *et al.* 2014; Webster and Lusk 2016; Webster *et al.* 2016). After replacing GFP_11_-mCherry-Pus1 with GFP_11_-Pus1, we examined colocalization of split-GFP signal in Pom33-GFP_1-10_ strains using mCherry-tagged proteins marking each of these NE puncta-associated structures. In most cells, Pom33-GFP_1-10_ did not colocalize with any of the tested NE proteins (Figure S4). Moreover, if we deleted key components involved in the formation of nuclear sub-complexes, we did not see an effect on the size or frequency of Pom33 puncta (Figure S4; data not shown). This result was particularly surprising as it suggests that Pom33 puncta represent a novel nuclear sub-compartment that is distinct, particularly from the SINC that has been previously described for incompletely assembled NPCs (Webster *et al.* 2014; Webster *et al.* 2016).

To better understand the nature of the Pom33 puncta, we were interested in determining if soluble nucleoporins not present in our screen were also present in foci. We localized at least one component from each NPC subcomplex in wild-type and *asi1∆* cells, including the outer ring components Nup120 and Nup145C, the inner ring protein Nup188, the central channel component Nup57 and the basket proteins Nup2 and Mlp2 (Figure 5A) (Aitchison and Rout 2012; Alber *et al.* 2007). Nup145C, Nup57, Nup2 and Mlp2 were unaffected by removal of *ASI1*; however, Nup120 and Nup188 exhibited punctate foci specifically in *asi1∆* mutants (Figure 5B). Over half of the Nup120-mCherry or Nup188-mCherry puncta seen in *asi1∆* colocalize with Pom33-GFP_1-10_ (Figure 5C). While it is tempting to speculate that these puncta contain full or complete NPCs, it is important to note that we only observe puncta formation with a few nucleoporins. While we detected Pom33 in *asi1∆* by immuno-electron microscopy (EM) at NPCs, there was no evidence of NPC clustering. Further, Pom33 did not localize to any recognizable NE landmark other than NPCs (data not shown), pointing to the possibility that Pom33 redistributes to a novel sub-nuclear component in the absence of Asi1.

**Figure 5.**
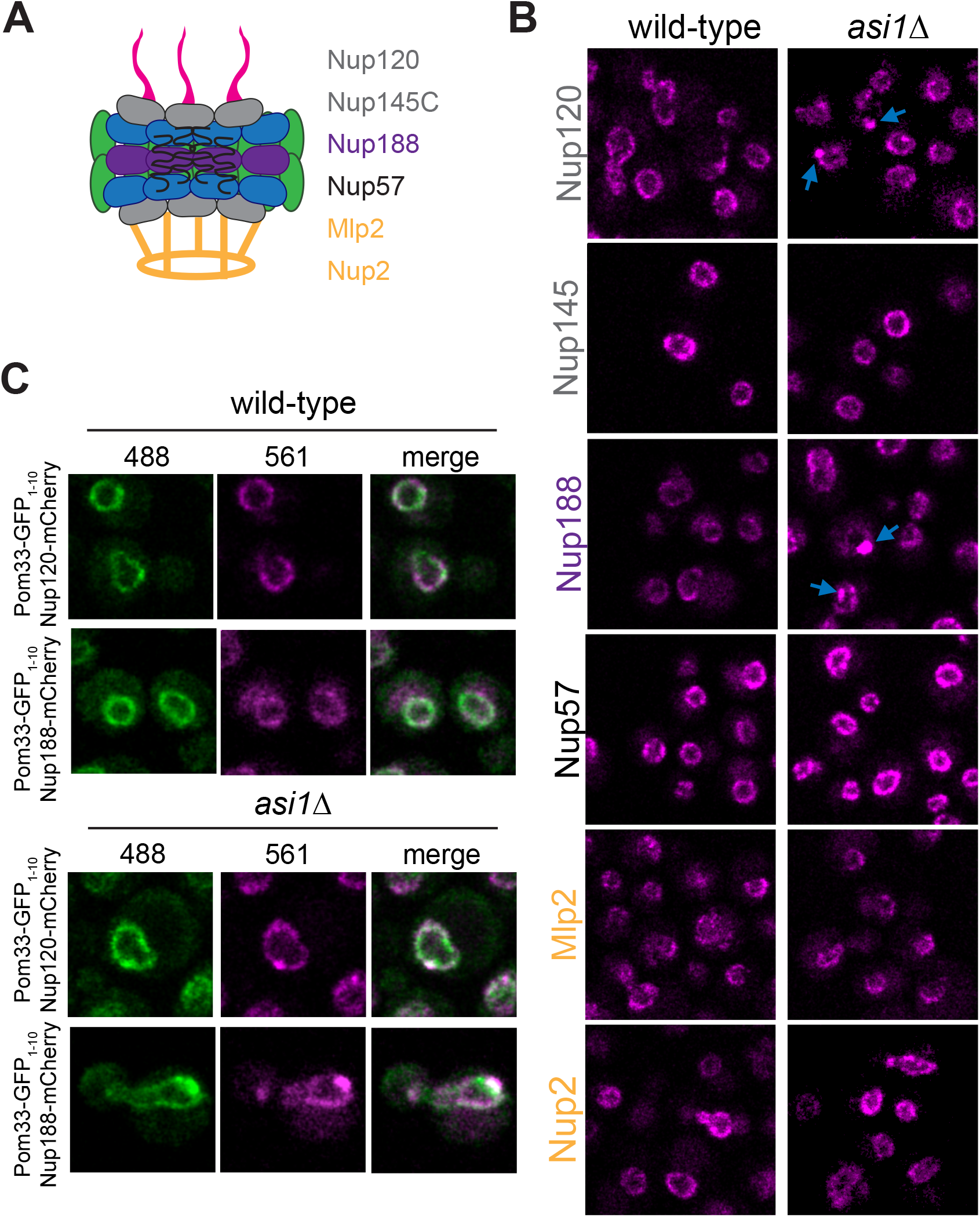
Nup188 and Nup120 colocalization with Pom33 puncta in. *asi1∆*. A. Cartoon of the NPC. Nucleoporins tested included the outer ring components Nup120 and Nup145C, the inner ring Nup188, the central channel nucleoporin Nup57 and the basket associated Mlp2 and Nup2. B. Localization of nucleoporins in A were assayed in wild-type and *asi1∆*. Nup120 and Nup188 displayed more puncta in *asi1∆,* as indicated by blue arrows. C. Example images of Pom33-GFP_1-10_ with either Nup120-mCherry or Nup188-mCherry in wild-type and *asi1∆*. Bar, 2 μm.

## Discussion

Mechanisms that control INM protein dynamics are poorly understood. Using split-GFP, we performed a systematic screen of over 700 membrane proteins, comparing INM levels between wild-type cells and mutants in the E3 ubiquitin ligases Asi1, Doa10 and Hrd1, the E2 ubiquitin conjugating enzyme Ubc7 and the vacuolar peptidase Pep4. Our data overlaps to a limited degree with previous studies examining substrates of these enzymes (Foresti *et al.* 2014; Khmelinskii *et al.* 2014). However, our ability to specifically and unequivocally assay changes at the INM enabled us to extend our understanding of INM quality control in several important ways. Previous work hypothesized that the primary role of INMAD is to remove ‘foreign’ substrates from the INM—proteins that have diffused in via NPCs but failed to diffuse back out, due to associated protein tags or lesions within the polypeptide (Foresti *et al.* 2014; Khmelinskii *et al.* 2014). However, we found that deletion of *ASI1* preferentially affected INM components compared to other mutants examined. In both our screen and our follow-up studies, we show that loss of *ASI1* results in increased levels of bonafide INM proteins, such as Pom33, Per33, Asi2 and Heh2. This suggests that Asi1, and by extension INMAD, plays a role in proteostasis of wild-type INM proteins, not just proteins mistargeted to the INM.

The effects of *ASI1* deletion on Pom33 distribution also shed light into the INMAD pathway. Asi1 and Asi3 are thought to form a heterodimeric E3 ligase (Foresti *et al.* 2014; Zargari *et al.* 2007), therefore, we anticipated that deletion of *ASI3* would mimic *asi1∆*. However, we found that the number of Pom33 foci in *asi3∆* was not statistically distinguishable from wild-type whereas it more than doubled in *asi1∆* (Figure 2). A similar observation was also made during studies of Asi1 regulation (Pantazopoulou *et al.* 2016), pointing to an incomplete understanding of the core INMAD enzymatic machinery. Given that Asi1 and Asi3 are multi-spanning membrane proteins, it has been difficult to set up in vitro biochemical assays to study their activity. However, several lines of evidence suggest that Pom33 is a direct Asi1 substrate. First, Pom33 is ubiquitinated in an Asi1-dependent manner. Second, expression of Asi1-DUb recapitulated the Pom33 puncta phenotype seen in *asi1*∆ mutants. Lastly, a constitutively ubiquitinated Pom33 (Ub-Pom33) largely reversed puncta formation in *asi1∆*. Taken together, these data suggest that Asi1 normally acts to ubiquitinate a pool of Pom33, which results in its uniform NE distribution.

Asi1-dependent Pom33 ubiquitination does not appear to be a signal for degradation. A ladder of poly-ubiquitinated Pom33 was not observed. In addition, Pom33 levels did not change when it was fused to ubiquitin or expressed in a proteasome mutant. Our data suggest that ubiquitination regulates Pom33 localization in much the same way that mono-ubiquitination is used throughout the endomembrane system to control subcellular localization or endocytosis (D’azzo *et al.* 2005; Hicke and Dunn 2003). Analysis of Pom33 localization determinants in budding yeast suggest that the C-terminal 65 amino acids are important for both stability and NPC localization (Floch *et al.* 2015), making it tempting to speculate that residues in this region are ubiquitinated. Attempts to map a ubiquitinated lysine residue were unsuccessful, suggesting that multiple lysines are sufficient (data not shown). Examination of our Asi1 substrates, along with previously described targets, did not show a particular motif targeting them to the ligase. However, it has been previously proposed that amphipathic helices play a role in Asi2 degradation and Doa10 substrate recognition (Boban *et al.* 2014; Ravid *et al.* 2006). Given that Pom33 has at least two amphipathic helices in its C-terminus that play a role in NPC targeting and membrane binding, one hypothesis is that these domains are also important for Asi1-dependent ubiquitination. More generally, INMAD-dependent control of these common motifs could play a role in regulation of NPC assembly and NE compartmentalization.

What is the role of INMAD regulation of Pom33 and other resident INM proteins? No obvious changes to nuclear structure, including NPC distribution, have been reported for *asi1∆* by EM (Foresti *et al.* 2014). Colocalization with Nup120 and Nup188 suggests that the Pom33 puncta contain at least some additional NPC components even though no recognizable NPCs were seen cytologically (data not shown). We do not believe that these are improperly assembled NPCs, as puncta did not colocalize with the SINC. Instead, our data suggests that Pom33 aggregates in a novel INM structure. Since the INMAD has only been investigated for its role in degradation of targets, it will be of future interest to determine whether the INMAD components mediate ubiquitination of other proteins to alter their localization or if TMEM33, the metazoan ortholog of Pom33, is also ubiquitinated.

## Materials & Methods

### Yeast strains and plasmids

All strains are derivatives of BY (*can1Δ::STE2pr-Sp-HIS5 lyp1Δ his3Δ1 leu2ΔO ura3ΔO met15ΔO LYS2*). The split-GFP library was made by first integrating the pRS315-*NOP1pr-GFP_11_-mCherry-PUS1* (pSJ1321) reporter into a MATα derivative to create SLJ7859 (MATα *can1Δ::STE2pr-Sp-HIS5 lyp1Δ his3Δ1 leu2ΔO ura3ΔO met15ΔO LYS2* p*LEU2*-*NOP1pr-GFP_11_-mCherry-PUS1*). Genes were C-terminally tagged with *GFP_1-10_-URA3MX* (pSJ1256) by PCR as previously described. These 1010 strains were crossed to deletion mutants taken from the yeast deletion collection (MAT*a yfg∆::KANMX LYP1 CAN1 his3 ura3 leu2*) using the Singer RoToR robot as previously described (Tong and Boone 2006). Following diploid selection on SD/MSG-Ura+G418, cells were sporulated for 3-4 weeks then haploids containing the deletion and the tags were selected twice on SD/MSG-Ura-His-Lys-Arg+thialysine+canavanine+G418. All plates were incubated at 23°C. We did not recover certain combinations of deletion mutants/tagged genes/selection markers due to linkage and slow growth of certain strains. Therefore, mutant libraries contained fewer genes than the original library. A complete list of genes screened can be found in Tables S1 and S2.

For colocalization experiments, we used the pRS315-*NOP1pr-GFP_11_-PUS1* (pSJ1679) Reporter so that genes could be C-terminally tagged with mCherry using *pFA6-mCherry-KANMX* by PCR. Strains used for each experiment are listed in Table S4, with the exception of strains taken directly from our split-GFP libraries.

### INM mutant screen

Cells were grown overnight at 23°C in SC-Leu in 96 well plate format, using deep well dishes (Thermo-Scientific). Imaging plates (Ibidi) were pretreated with 100 μl of poly-lysine (Sigma-Aldrich) for 1 h and then rinsed twice with water and allowed to dry before use. Cells were plated using 100 μl of culture. Each plate was imaged on a Nikon Eclipse TI equipped with a Yokogawa CSU W1 spinning disk head and Andor EMCCD using a Nikon Apo TIRF 100X 1.49NA Oil objective. mCherry was imaged using a 561nm laser at 100% power and ET605/70m emission filter, with an exposure time of 100 ms. GFP was imaged using a 488nm laser at 100% power and ET535/30m emission filter, with an exposure time of 200 ms. Four points were automatically selected for each well. An automation script moved to positions, found focus using Nikon PFS, and imaged each channel with a z-stack of 13 slices and spacing 0.5 μm. Image processing was performed in ImageJ using custom macros and plugins. In brief, images were background subtracted and sum projected in z, and for each cell a mask of the nucleus was generated based on the mCherry channel, as well as a ring mask surrounding the nucleus to measure cytoplasmic signal. mCherry and GFP intensity was measured in the nuclear and ring masks for each cell. Cells with high cytoplasmic signal were discarded as dead cells. For each live cell, the nuclear GFP/mCherry intensity ratio was calculated; from these data the average ratio per sample was calculated, and for each protein the mutant/wild-type ratio was calculated from these averages (given in Table S1). Proteins were counted as hits if the mutant/wild-type ratio was greater than 1.4.

Wild-type and *asi1∆* images were manually inspected for signal in both the 561 nm and 488 nm channels. If no signal was present upon visual inspection, the sample was categorized as a false-positive and was removed from subsequent analysis with all mutants. These genes are listed in Table S2; most correspond to negatives or non-abundant INM hits in wild-type cells (Smoyer *et al.* 2016).

### Confocal imaging

Cells were grown overnight in SC-Leu in 2 ml cultures and then harvested for imaging and immobilized on poly-lysine slides (Polysciences Inc). Imaging was conducted on a Zeiss-LSM780. All images were taken using a 40x water objective, fluorophores were excited using a 488-nm and 561 nm Argon laser line, GFP emission was collected through a BP 505-540 nm filter and mCherry was collected through a LP 580 nm filter, with the pinhole set to 1 airy units. Images taken for puncta quantification, puncta measurement and colocalization experiments had a zoom of 6, z stack of 10-12 slices. Puncta widths were measured from manually drawn line profiles over puncta averaged over a thickness of 8 pixels. Profiles were then aligned to their peak maxima and averaged to determine average peak profiles. They were then fit to Gaussian functions using non-linear least squares to determine their width. Error bars were determined from monte carlo analysis with 100 random simulations as previously described (Bevington and Robinson 2003).

### Ubiquitination of Pom33

Liquid nitrogen ground lysates were prepared as previously described in (Bupp *et al.* 2007; Friederichs *et al.* 2012; Jaspersen *et al.* 2006). Cells of indicated genotypes were grown overnight, then diluted into 1 liter cultures in the morning to an OD_600_ of ~0.2. Cells were allowed to grow until mid-log phase and then harvested. Extraction buffer was adjusted for each sample so that an OD_600_ of 1.0 would have 5 ml of extraction buffer added. Cells were then dropped into liquid nitrogen to make yeast dots and then frozen at -80°C. Yeast powder was made by grinding the frozen yeast dots using Retsch ball mill 4 x 3 min, 30 Hz. Powder was then collected and weighed out to equal amounts for each sample. Ground cell powder was thawed on ice, then resuspended in 9 mL of extraction buffer (20 mM Hepes-NaOH, pH 7.5; 300 mM NaCl; 1 mM EDTA; 5 mM EGTA; 50mM NaF; 50 mM β-glycerophosphate; 0.5% TritonX-100; 1 mM DTT; 1 mM PMSF; and 1 mg/mL each pepstatin A, aprotinin, and leupeptin). Next, samples were homogenized with a Polytron 10/35 for 30 sec, lysates were centrifuged at 3000 × g for 10 min. at 4°C and the resulting supernatant was used for immunoprecipitations. Lysates were added to protein G Sepharose beads (GE Healthcare) that had been previously incubated with P4D1 anti-Ubiquitin antibody (1:500 μl, Cell Signaling Technology) and rotated for ~ 3 hours at 4°C. Beads were washed 5x with extraction buffer before western blot analysis, using (1:1000) anti-GFP antibody (Cell Signaling Technology).

### Cycloheximide chase experiments

Cells were grown overnight and diluted back the next morning, then allowed to recover to mid-log phase before the addition of cycloheximide (125 μg/ml). Samples were removed at given time points and quick frozen in liquid nitrogen for further analysis by western blotting, using (1:1000) anti-MYC antibody (Cell Signaling Technology), (1:10000) anti-histone H4 antibody (Abcam) and (1:1000) anti-Clb2. For the *cim3-1* experiments, cells were shifted to the non-permissive temperature of 37°C for 45 min before the addition of cycloheximide.

## Acknowledgements

We are grateful to Chris MacDonald and Robert Piper for the Asi1-DUb construct. We are grateful to Richard Alexander, Sean McKinney, Brian Slaughter and Melainia McClain for help during this project and to Brian Slaughter and members of the Jaspersen lab for comments on the manuscript. SLJ is supported by the Stowers Institute for Medical Research. CJS is a predoctoral researcher in the Graduate School of the Stowers Institute. Original data underlying this manuscript can be downloaded from the Stowers Original Data Repository at http://www.stowers.org/pubs/LIBPB-1363.

## Author Contributions

CJS and SLJ conceived of applying split-GFP to study degradation of INM proteins, SM made the yeast libraries that were screened by CJS with assistance from SES. Intensity ratios were measured and analyzed by CJS and SES using image analysis tools developed by JRU and SES. CJS and SLJ wrote the paper with input from all the authors.

## Abbreviations

NPC: nuclear pore complex
INM: inner nuclear membrane
ONM: outer nuclear membrane
NE: nuclear envelope
ER: endoplasmic reticulum
INMAD: inner nuclear membrane-associated degradation
ERAD: ER-associated degradation
Ub: ubiquitin
DUb: deubiquitin
SGD: Saccharomyces cerevisiae genome database
GO: Gene Ontology
MW: molecular weight
FWHM: full width half maximum
SPB: spindle pole body
EM: electron microscopy
NVJ: nuclear-vacuolar junction
INQ: intranuclear quality control
SINC: storage for incomplete NPCs

---

**Figure S1.**
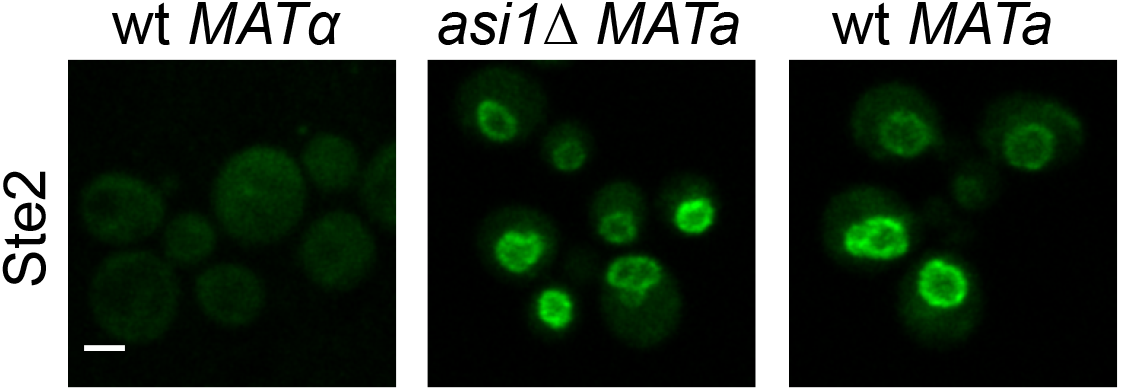
Ste2 localization in Mata and Matα. Ste2-GFP_1-10_ INM signal is mating-type specific. Bar, 2 μm.

**Figure S2.**
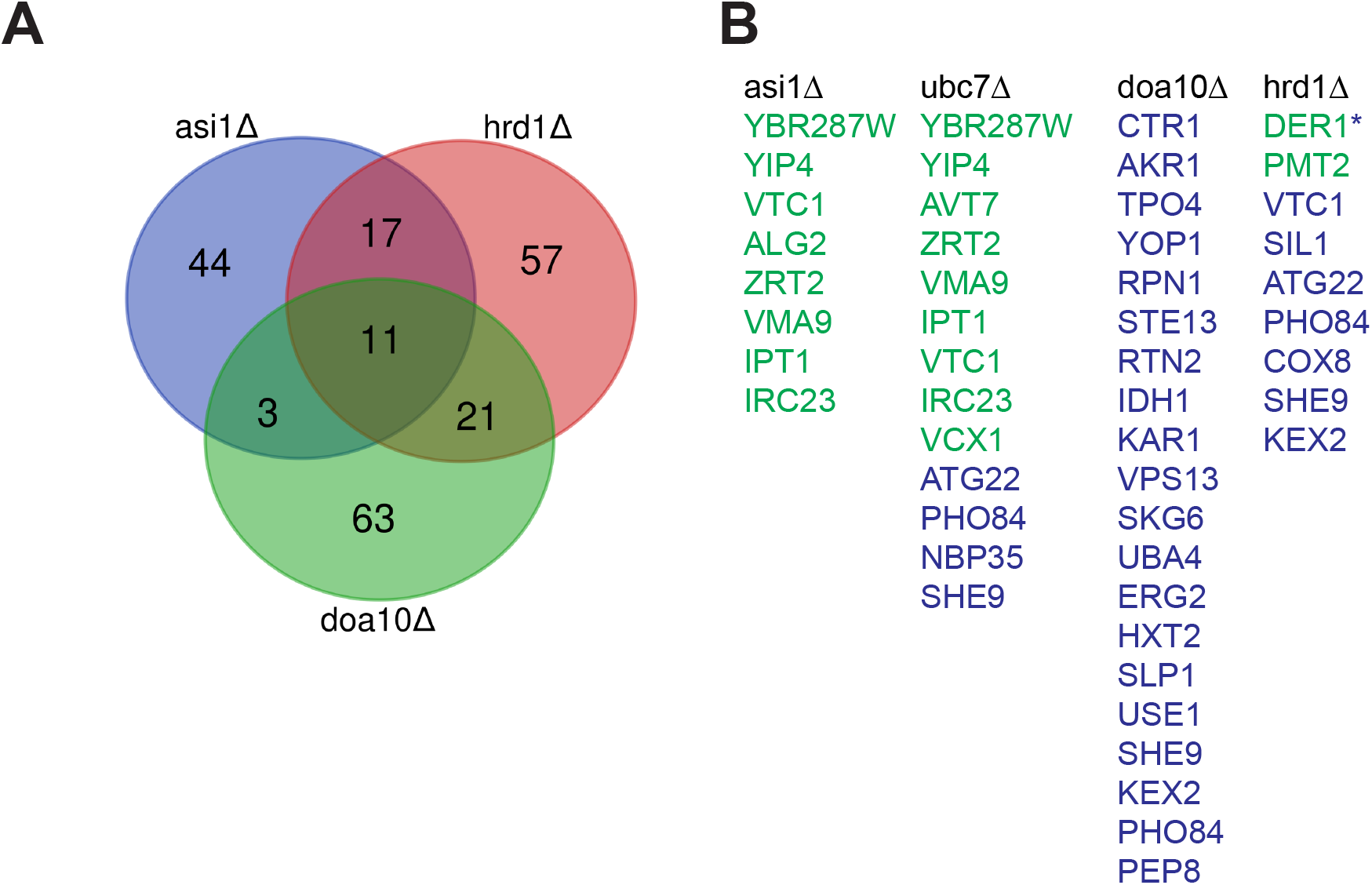
Overlap of potential targets within our screen and previously reported targets. A. Venn diagram showing overlap of hits for the E3 ligase mutants *asi1∆*, *doa10∆* and *hrd1∆*. B. Comparison of targets identified in our screen with those reported for the tandem fluorescent timer screen (Khmelinskii et al. 2014) in green and proteomic analysis (Foresti et al. 2014) in blue. An asterisk denotes targets common in all three reports.

**Figure S3.**
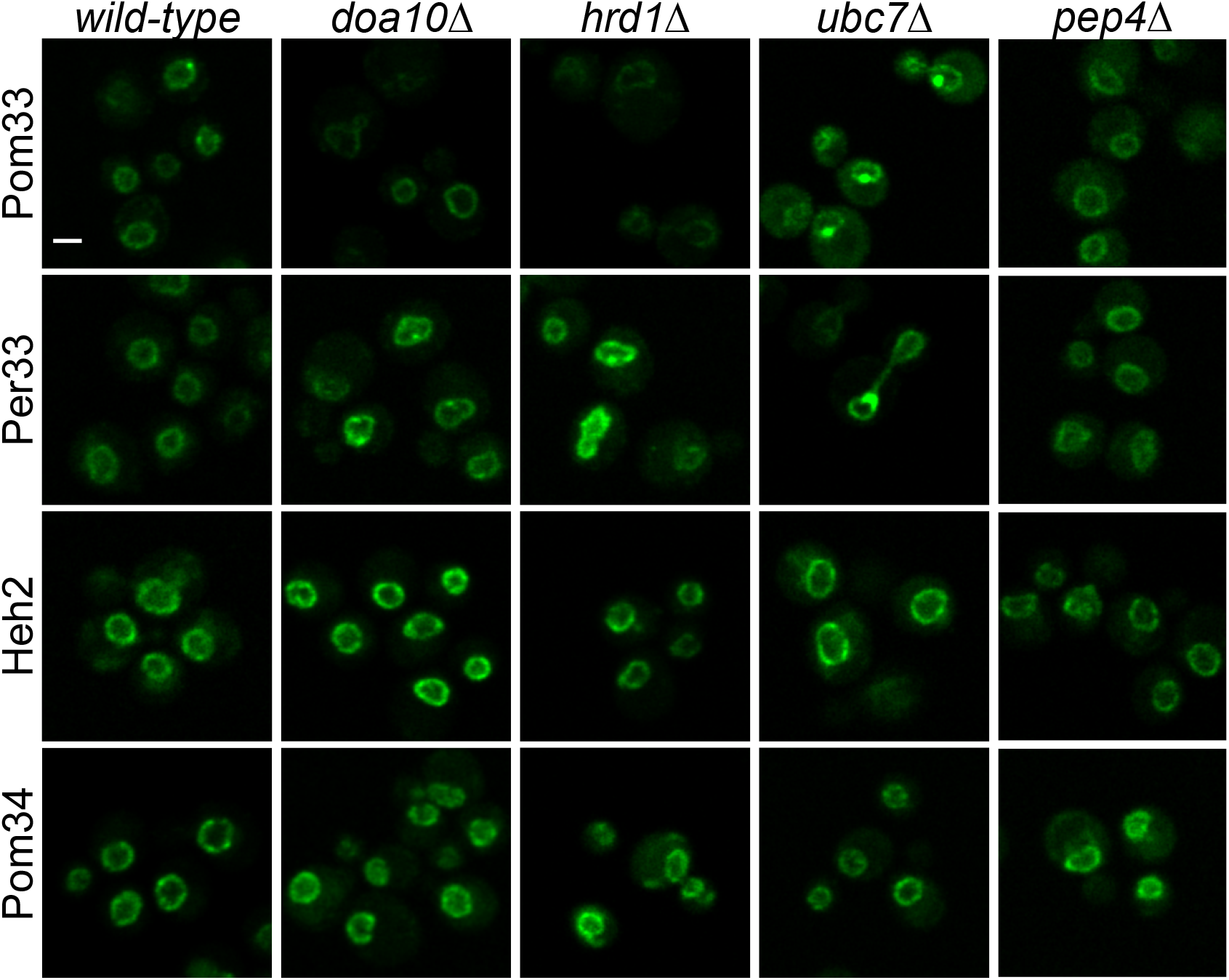
Puncta are specific to Pom33 in mutants of the Asi Complex and E2 enzyme Ubc7. Example images of Pom33, Per33, Heh2, Pom34 signal in mutants disrupting the membrane E3 ligases Doa10 and Hrd1, a mutant of the E2 enzyme, Ubc7 and a mutant of the vacuolar protease, Pep4. Bar, 2 μm.

**Figure S4.**
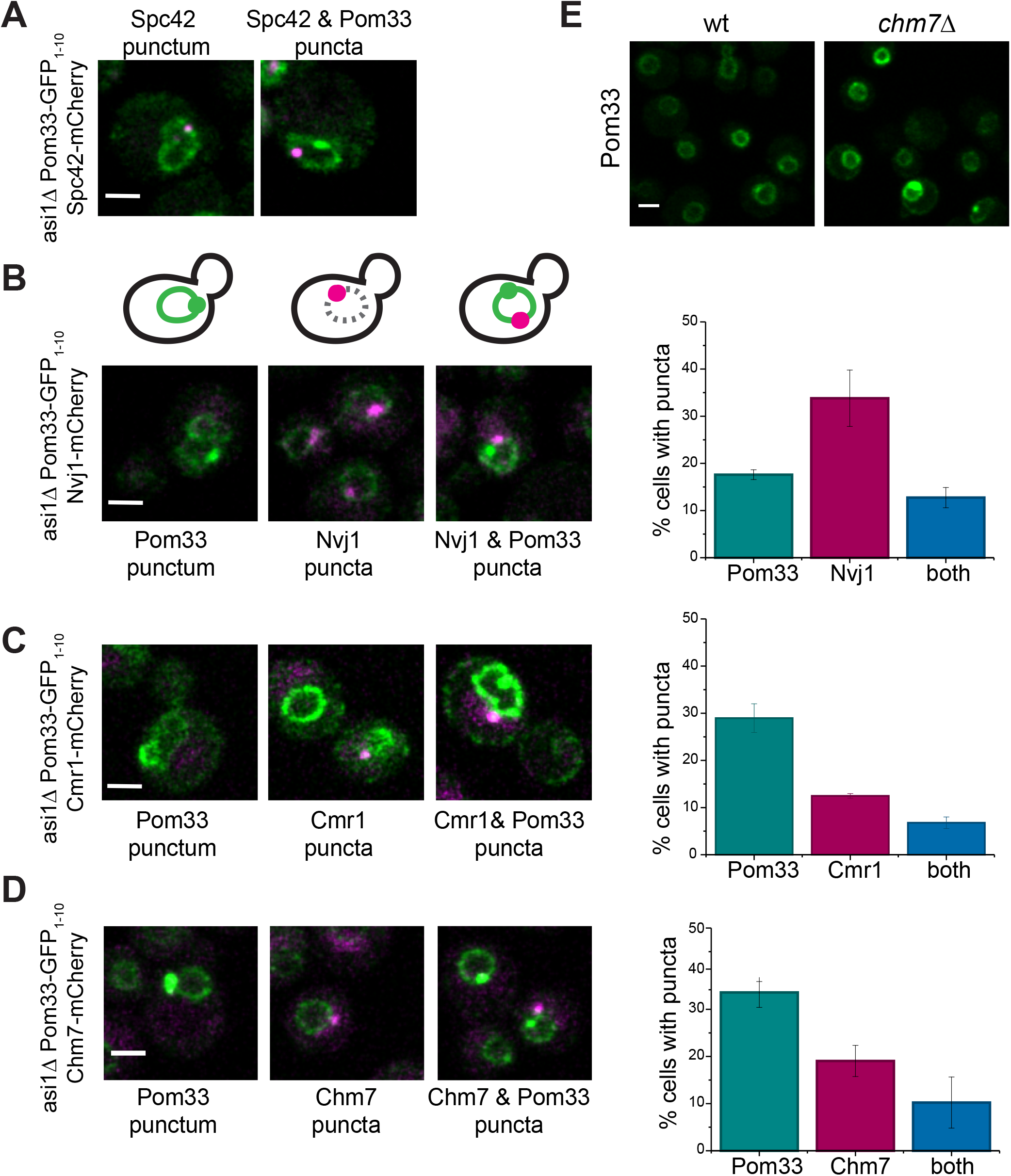
Pom33 puncta tested with known nuclear foci representing the SPB, NVJ, INQ and SINC. A. Pom33-GFP_1-10_ puncta did not colocalize with the SPB. B. Left, example images of Pom33-GFP_1-10_ puncta, Nvj1-mCherry puncta (NVJ), and both in the same cell. Right, quantitation of puncta frequency, n = 198 cells total. C. Left, example images of Pom33-GFP_1-10_ puncta, Cmr1-mCherry puncta (INQ), and both in the same cell. Right, quantitation of puncta frequency, n = 80 cells total. D. Left, example images of Pom33-GFP_1-10_ puncta, Chm7-mCherry puncta (SINC), and both in the same cell. Right, quantitation of puncta frequency, n = 150 cells total. E. Disruption of *CHM7* did not change Pom33 puncta formation. Bars, 2 μm. Error bars equal SEM.

